# ATP-Independent Nucleosome Unfolding by FACT: Electron Microscopy Analysis

**DOI:** 10.1101/2021.07.13.452273

**Authors:** Anastasiia L. Sivkina, Maria G. Karlova, Maria E. Valieva, Laura L. McCullough, Timothy Formosa, Alexey K. Shaytan, Alexey V. Feofanov, Mikhail P. Kirpichnikov, Olga S. Sokolova, Vasily M. Studitsky

## Abstract

FACT is a histone chaperone that unfolds nucleosomes without ATP hydrolysis. We used electron microscopy to study FACT and FACT:nucleosome complexes, and found that both adopt broad ranges of configurations, indicating high flexibility. We found unexpectedly that the DNA binding protein Nhp6 also binds to the C-terminal tails of FACT subunits, inducing more open geometries of FACT even in the absence of nucleosomes. Nhp6 therefore supports nucleosome unfolding by altering both FACT structure and nucleosome properties. Complexes formed with FACT, Nhp6, and nucleosomes also produced a broad range of structures, revealing a large number of potential intermediates along a proposed unfolding pathway. The data suggest that Nhp6 has multiple roles before and during nucleosome unfolding by FACT, and that the process proceeds through a series of energetically similar intermediate structures, ultimately leading to an extensively unfolded form.

**One Sentence Summary:** Electron microscopy reveals the pathway of ATP-Independent nucleosome unfolding by histone chaperone FACT.

## Main Text

The eukaryotic genome is densely packed into nucleosomes, each containing 145-147 bp of DNA^1,2^. This packing blocks the accessibility of the DNA to many of the proteins that control gene expression, with access tightly regulated by many factors including ATP-dependent remodelers and ATP-independent histone chaperones^3–6^. FACT (facilitates chromatin transcription) is a broadly conserved histone chaperone that promotes both large-scale nucleosome unfolding and nucleosome assembly, contributing to multiple phases of transcription, replication, and repair^7,8,6^.

The larger Spt16 (Suppressor of Ty) subunit of FACT is similar in all eukaryotes, while the smaller subunit has two variants; the Pob3 (Polymerase One Binding) version found in yeasts and the SSRP1 (Single-strand Recognition Protein) found in higher organisms^9^. The primary difference is that SSRP1 includes an HMGB-family DNA-binding domain that is absent in Pob3. Yeast FACT activity is enhanced both *in vitro* and *in vivo* by the HMGB-domain factor Nhp6 (Non-Histone Protein), but Nhp6 can also drive the activity of human FACT, so the functions of the HMGB domains and the reason for the distinct architectures of Pob3 and SSRP1 are unknown^10^. Spt16 and Pob3/SSRP1 are organized into multiple, flexibly-associated structural domains that contain several binding sites for H3/H4 tetramers and H2A/H2B dimers, so FACT can interact simultaneously with all of the components of nucleosomes (see ^4–6^ for review).

The nucleosome unfolding and assembly activities of FACT are thought to function in different physiological processes, with unfolding participating in efficient removal of nucleosomes from promoters during induction of transcription^11,12^, and assembly or stabilization of nucleosomes being more important for nucleosome survival during transcription^13^ and as chromatin is deposited during repression and replication^14–16^. Notably, unfolding activity is conserved between yeast and human FACT, but requires Nhp6 in both cases^7,10,17^, and can also be supported by the small molecule DNA intercalators known as curaxins^18^.

The structures of individual domains of FACT have been revealed by crystallography^4^, and a recent cryo-EM structure provided a view of how these domains can collaborate to destabilize nucleosomes^19^. However, this structure was based on an unusually stable complex of FACT with a nucleosome lacking entry/exit point DNA, artificially exposing the binding site for the C-terminal tail of Spt16 on the H2A-H2B surface. The structure did not reveal the locations of the N-terminal domain of Spt16 or the HMBG and C-terminal domains of SSRP1, and the DNA remained coiled^19^, unlike its status in the fully unfolded form^7^. The available structure therefore did not resolve how the HMGB domain contributes to nucleosome unfolding and it appears to represent just one of many potential steps along the pathway to the unfolded state.

Here we report our analysis of FACT, FACT:Nhp6 and FACT:Nhp6:nucleosome structures by transmission electron microscopy and single particle FRET (spFRET). We observed a range of structures, consistent with the association of the domains of FACT with one another through flexible linkers, but we were able to group these into subsets by 2D class averaging, suggesting favored conformations for both FACT alone and for FACT:nucleosome complexes. Importantly, we found that Nhp6 binds to the acidic C-terminal tails of both Spt16 and Pob3, altering the distribution of configurations in FACT before it binds to nucleosomes, and supporting the reversible unfolding of nucleosomes to a nearly linear structure. We were also able to arrange the populations of averaged structures into a proposed pathway for unfolding, revealing a potential series of sequential steps in this process.

## Results

### Nhp6 protein interacts with the acidic C-terminal domains of Spt16/Pob3

Human FACT has a single HMGB domain within the SSRP1 subunit, but its function is supplied by the separate Nhp6 protein in yeast (**Fig. 1a** and ^7,10,17^). Some Nhp6 immunoprecipitated with Spt16/Pob3 heterodimers from whole cell lysates prepared at low ionic strength, but Nhp6 does not co-purify with Spt16/Pob3 under physiological conditions so it is not a stable, stoichiometric subunit of FACT^9,20^. We reinvestigated this association by non-denaturing gel electrophoresis, and found that Nhp6 reduced the migration of Spt16/Pob3, suggesting at least transient formation of complexes (**Fig. 1b**).

**Figure 1.**
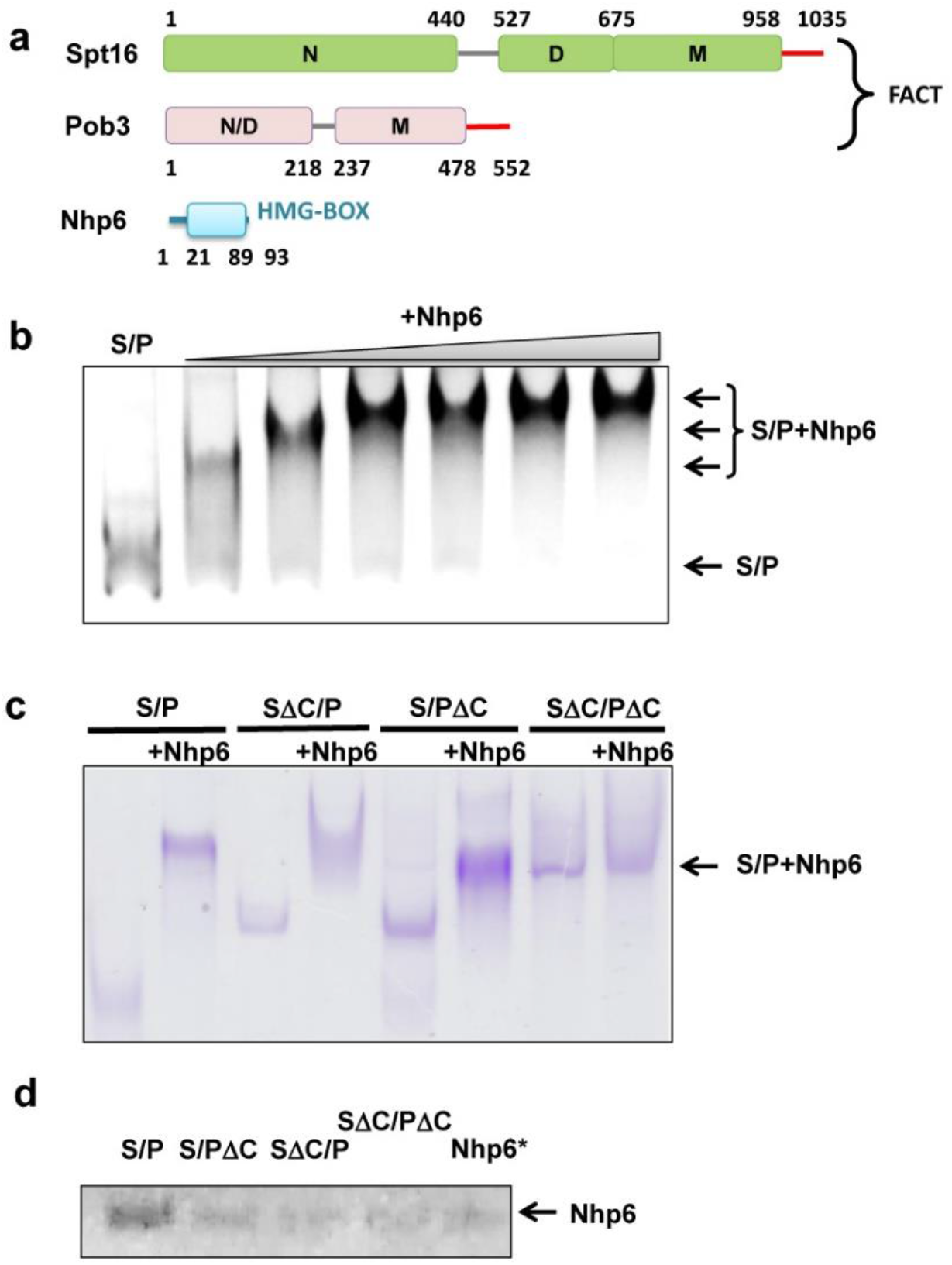
Nhp6 protein interacts with C-terminal domains of FACT subunits. **a.** FACT and Nhp6 domain structures. FACT is a dimer of Spt16 and Pob3 subunits and requires Nhp6 protein for nucleosome unfolding. N – N-terminal domain; D – dimerization domains; M – middle domains; N/D – N-terminal/dimerization domain. Negatively charged C-terminal regions of Spt16 and Pob3 are shown in red. **b.** Spt16/Pob3 (S/P, 0.13 μM) was incubated with Nhp6 (0, 0.26 μM, 0.52 μM, 0.78 μM, 1.04 μM, 1.3 μM, or 2.6 μM) and analyzed by native PAGE followed by silver staining. Arrows indicate distinct migration patterns. **c**. Native PAGE analysis of the migration of FACT mutants lacking the C-terminal regions of Spt16 (SΔC), Pob3 (PΔC), or both, with or without Nhp6, stained with Coomassie blue. The arrow indicates the region excised to test for Nhp6. **d.** Bands (as in 1c) containing apparent FACT:Nhp6 complexes were excised, subjected to denaturing SDS-PAGE and silver stained. The region of the gel containing Nhp6 protein is shown. Nhp6* shows Nhp6 level in the empty area of the gel from the lane containing Nhp6 only, indicating the background level of Nhp6 detection.

HMGB-domain factors are basic proteins that bind to and bend DNA^21^, and titration of FACT with Nhp6 produces a pattern of migration similar to the one shown in **Fig. 1b**, with multiple intermediates corresponding to gradual saturation of binding sites. We considered the possibility that the acidic C-terminal regions found in both Spt16 and Pob3 provided multiple potential interaction sites for Nhp6. Consistent with this, deleting either C-terminal domain altered the electrophoretic mobility shift induced by Nhp6, and deleting both domains essentially eliminated the effect (**Fig. 1c**). FACT heterodimers lacking these domains were stable but were inactive in a reorganization assay^22^; these same regions contain the primary binding sites for H2A/H2B dimers. Our results therefore show that FACT uses the same C-terminal domains to interact with H2A/H2B and with Nhp6.

To determine whether the apparent complexes contain Nhp6, these regions of the native gel were excised and subjected to SDS-PAGE followed by silver-staining. As shown in **Fig. 1d**, a small amount of Nhp6 was detected in a control region, but significantly more was observed in the region containing full-length Spt16/Pob3. As reported previously, Nhp6 migrates in a broad band in native gels^9^, so it is difficult to determine the stoichiometry of the complexes, but these results show that the impaired migration of Spt16/Pob3 in the presence of Nhp6 is largely due to the acidic C-terminal tails of both subunits of FACT, supporting a direct interaction between them and Nhp6.

### FACT is a flexible complex that is unfolded by Nhp6

FACT is composed of multiple globular domains connected by flexible linkers (**Fig. 1a**). To determine the range of conformations it adopts, we examined Spt16/Pob3 alone or with Nhp6 by transmission EM at a magnification of 40,000x (**Fig. 2a**). Multiple images of FACT ± Nhp6 were obtained, yielding 10,304 FACT particles and 28,425 FACT:Nhp6 particles (**Supplementary Table S1**). These were analyzed by reference-free 2D classification. 112 classes (100-200 particles per class) with distinct features were identified for each type of complex (**Supplementary Fig. S1**).

**Figure 2.**
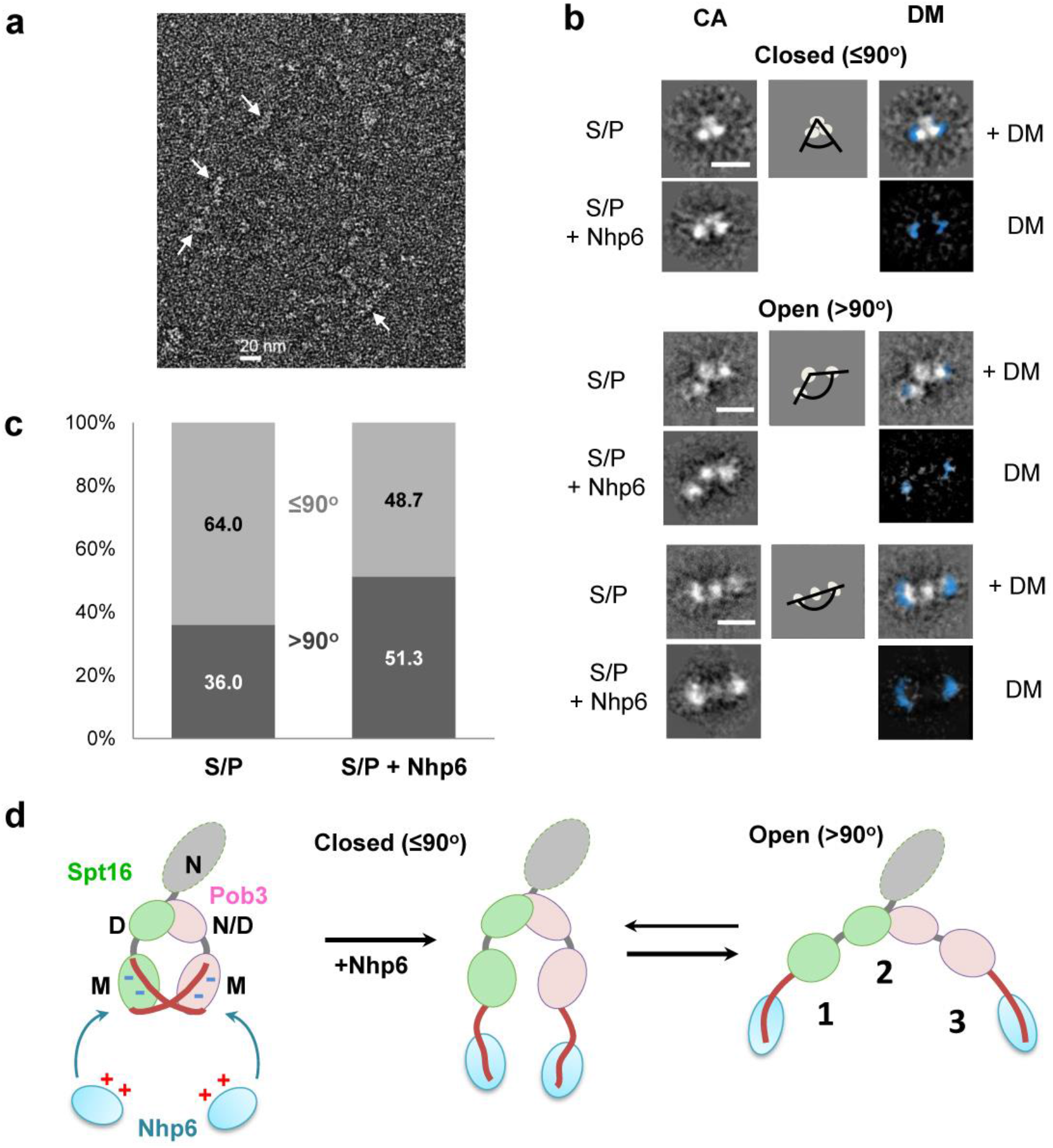
FACT is a flexible complex that is unfolded by Nhp6. **a.** Representative images of Spt16/Pob3 obtained by transmission electron microscopy after negative staining. Arrows indicate single FACT particles. **b.** Representative 2D class averages of Spt16/Pob3 with different arrangements of the three resolved densities in the presence/absence of Nhp6. Scale bar: 10 nm. CA – class average, DM – difference map. **c.** Quantitation of particles identified as closed (≤90°) or open (≥90°) in samples with and without Nhp6. **d.** Potential identities of the three resolved densities detected in class averages and a model for how Nhp6 promotes formation of a more open form are shown. The grey oval on the scheme is Spt16N domain which was not resolved.

**Fig. 2b** shows representative structural classes observed. The particles all contained three interconnected electron densities but multiple geometries were detected, consistent with flexible linkages (**Fig. 2b**, **Supplementary Movies S1, S2**). The classes fell generally into groups with more compact, “closed” conformations (~8.0 x 6.9 nm) and more linear, “open” forms (~13.1 x 4.2 nm) (**Fig. 2b**). Based on the structure of human FACT with a nucleosome^19^, the two flanking electron densities are likely to be the M-domains of Pob3 and Spt16, while the middle density is likely to be the dimerized Pob3-N/D:Spt16-D domains (**Fig. 2d**). The Spt16-N domain was not identified in either the previous study^19^ or in our class averages, suggesting that it adopts too many conformations to be visible after averaging.

The tripartite structure characteristic of FACT alone was also observed in samples containing Nhp6 (**Fig. 2b**), but the flanking densities appeared to get larger. To test this, we aligned the class-averages with similar geometries and calculated difference maps, confirming extra density in the distal regions of the structure corresponding to the surfaces we assigned as the M domains (**Fig. 2b**). This is consistent with the conclusion above that Nhp6 bound to the acidic C-terminal tails where they protruded from the M domains (**Fig. 1**).

Notably, addition of Nhp6 also increased the fraction of particles in the open conformation from 36% for FACT alone to 51% in the FACT:Nhp6 complexes (**Fig. 2c**). The EM data therefore support a model in which Nhp6 binds to the acidic tails of Spt16 and Pob3, and suggest that this releases the middle domains to adopt a more open geometry (**Fig. 2d**). We propose that the acidic tails of each subunit interact electrostatically with positively charged surfaces of the other subunit in the absence of Nhp6, constraining the geometry of the heterodimers (**Supplementary Fig. S2**).

### FACT and Nhp6 unfold nucleosomes into a nearly linear protein-DNA structure

To examine how FACT affects the structure of intact nucleosomes, we inserted fluorescent dyes into a 147-bp DNA fragment based on the Widom 603 positioning sequence^23^ and assembled mononucleosomes with recombinant yeast histone octamers. Cy3 and Cy5 were placed at positions 35 and 112 bp from the edge of the nucleosome, bringing them close enough in the canonical nucleosome structure to allow efficient Förster resonance energy transfer (FRET) between the dyes^7^ (**Fig. 3a**). These nucleosomes were then used to probe the effects of FACT and Nhp6 on DNA uncoiling detected by single-particle FRET as previously described^7^ (**Fig. 3c**) and by in-gel FRET (**Fig. 3b**), and also to observe structural changes by EM (**Fig. 3d, e**).

**Figure 3.**
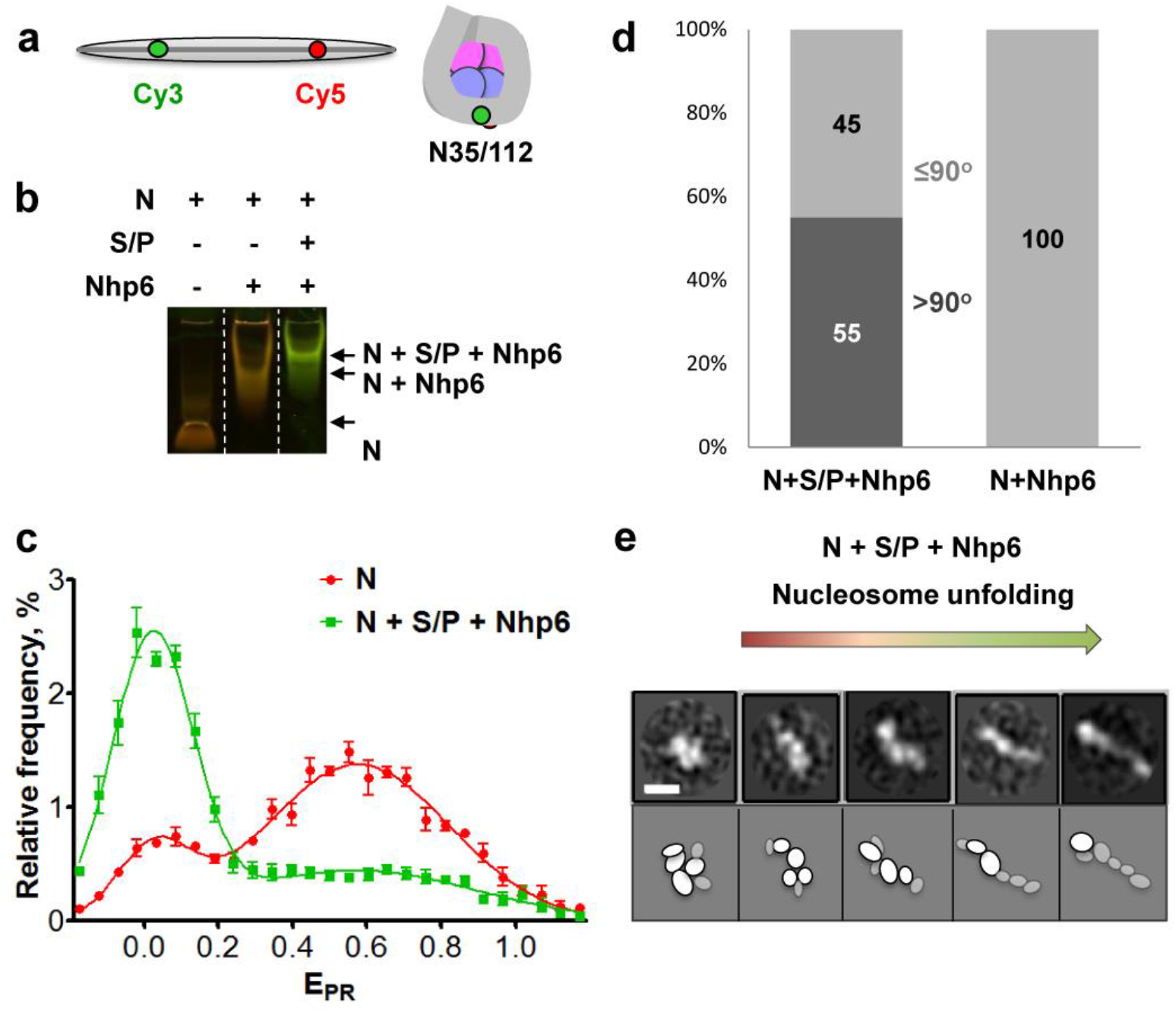
FACT and Nhp6 unfold nucleosomes into a nearly linear protein-DNA structure. **a.** Schematic of the Widom 603 sequence with Cy3 and Cy5 distant in the free DNA but adjacent in the N35/112 nucleosome (35 and 112 bp from the nucleosome boundary). **b.** Characterization of FACT:Nhp6 complexes with fluorescently labeled N35/112 nucleosomes (N) by in-gel FRET. Complexes were separated by native PAGE, and the gel was analyzed as described in Methods. Unfolding of nucleosomal DNA is detected by a decrease in FRET efficiency (transition from red/orange to green). **c.** Typical frequency distributions for nucleosomes N35/112 by the proximity ratios (E_PR_) in the absence (red curve) or in the presence of Nhp6 and FACT (green curve). E_PR_ profiles were calculated from the FRET efficiencies of individual nucleosomes in solution (average of three independent repeats, mean ± SEM). The maxima of E_PR_ peaks (mean ± SEM) were: N – 0.03 ± 0.00, 0.59 ± 0.03; (N+SP+Nhp6) – 0.02 ± 0.02, 0.55 ± 0.11. **d.** Fractions of closed (<90°) and open (>90°) complexes in FACT:Nhp6:nucleosome and Nhp6:nucleosome complexes. **e.** Representative 2D class averages of FACT:Nhp6:nucleosome complexes with different distances between edges of the complex are arranged to show the proposed sequence of events during nucleosome unfolding by FACT:Nhp6. Scale bar: 10 nm. Bottom: A schematic interpretation of the densities observed by EM to potential domains; less ordered densities are shown in grey. Note that the most compact complex shown on the left is similar to the FACT-subnucleosome complex described previously^19^.

We previously showed that (i) FACT induces large-scale uncoiling of nucleosomal DNA, separating DNA gyres carrying Cy3 and Cy5 dyes and reducing their FRET efficiency, (ii) this uncoiling requires high levels of Nhp6, and (iii) the uncoiling is reversible upon removal of FACT^7^. Uncoiling of nucleosomal DNA was also visible in native gels as a shift from orange (lower FRET) to green (higher FRET) color in FACT:Nhp6:nucleosome complexes, but not in Nhp6:nucleosome complexes (**Fig. 3b**). Nhp6 therefore binds to nucleosomes but does not induce uncoiling, consistent with previous results^7^. As further validation of nucleosomes, we characterized their structure by spFRET microscopy, which showed that 84.3±1.4% of the nucleosomes displayed high FRET (E_PR_ peak at 0.59 ± 0.03) and 15.7±1.4% had lower FRET (E_PR_ peak at 0.03 ± 0.00), revealing a distribution between canonical and uncoiled forms of nucleosomes in solution. As expected from our prior study^7^, FACT:Nhp6 increased the fraction of uncoiled forms to 68±5% (**Fig. 3c**).

Nhp6:nucleosome and FACT:Nhp6:nucleosome complexes were next isolated from native gels (**Fig. 3b**), transferred to hydrophilized copper grids, stained with 1% uranyl acetate, and analyzed by EM. Gel purification increased the fraction of particles in complexes, yielding particles that fell into 24 2D classes (**Supplementary Fig. S3**). While FACT alone produced three local densities in most images, complexes with nucleosomes typically contained 5-6 densities (**Fig. 3e** and **Supplementary Fig. S3**). Nhp6:nucleosomes were uniformly compact, but over half of the nucleosome complexes with FACT:Nhp6 displayed a more open conformation (**Fig. 3d**), similar to FACT:Nhp6 alone (**Fig. 2**), but with greater variation in the length of the particles (the distance between the lateral densities). The longest particles were nearly linear, suggesting a stepwise nucleosome unfolding pathway leading from a compact form to an extended one (**Fig. 3e**).

FACT:Nhp6:nucleosome classes had more densities than FACT:Nhp6, and they also had different patterns of distribution of the densities. These ranged from relatively compact conformations to the most elongated, thin shape that had a weak central density ~5 nm in width (**Fig. 3e**). Both compact and extended forms contained 5-6 densities, and both FACT:Nhp6 and FACT:Nhp6:nucleosome complexes had a similar fraction of open forms (51%-55%; **Fig. 2c**, **3d**). FACT:Nhp6:nucleosome complexes are therefore as flexible or more flexible than FACT:Nhp6 complexes, and while both can adopt compact and open forms, Nhp6 appears to drive the balance towards more open forms. The large number of configurations observed suggests that FACT distributes nucleosomes into many structural intermediates, not just canonical and unfolded forms.

### Mapping domains to observed densities

In order to assign the additional electron densities observed in FACT:Nhp6:nucleosome complexes to particular proteins/domains, we compared the structures of the open FACT:Nhp6 and FACT:Nhp6:nucleosome classes (**Fig. 4**). Even the most extended FACT:Nhp6:nucleosome complex was considerably shorter than a 147 bp DNA molecule, suggesting that ~70 bp of DNA was disordered and was not visualized (**Fig. 4a**). We propose that this disordered DNA extends from either side of the observed densities and could contain one or more molecules of Nhp6^16,33,34^. Based on the similarity of the central regions with and without nucleosomes, we propose that the Pob3-M, Pob3-N/D/Spt16-D, and Spt16-M domains make up the center of the complex with nucleosomes, with the histones and possibly additional Nhp6 molecules adding density and further lateral extension (**Fig. 4b**). As discussed above, the observed densities are less compact in the structures with nucleosomes, consistent with greater flexibility and less order.

**Figure 4.**
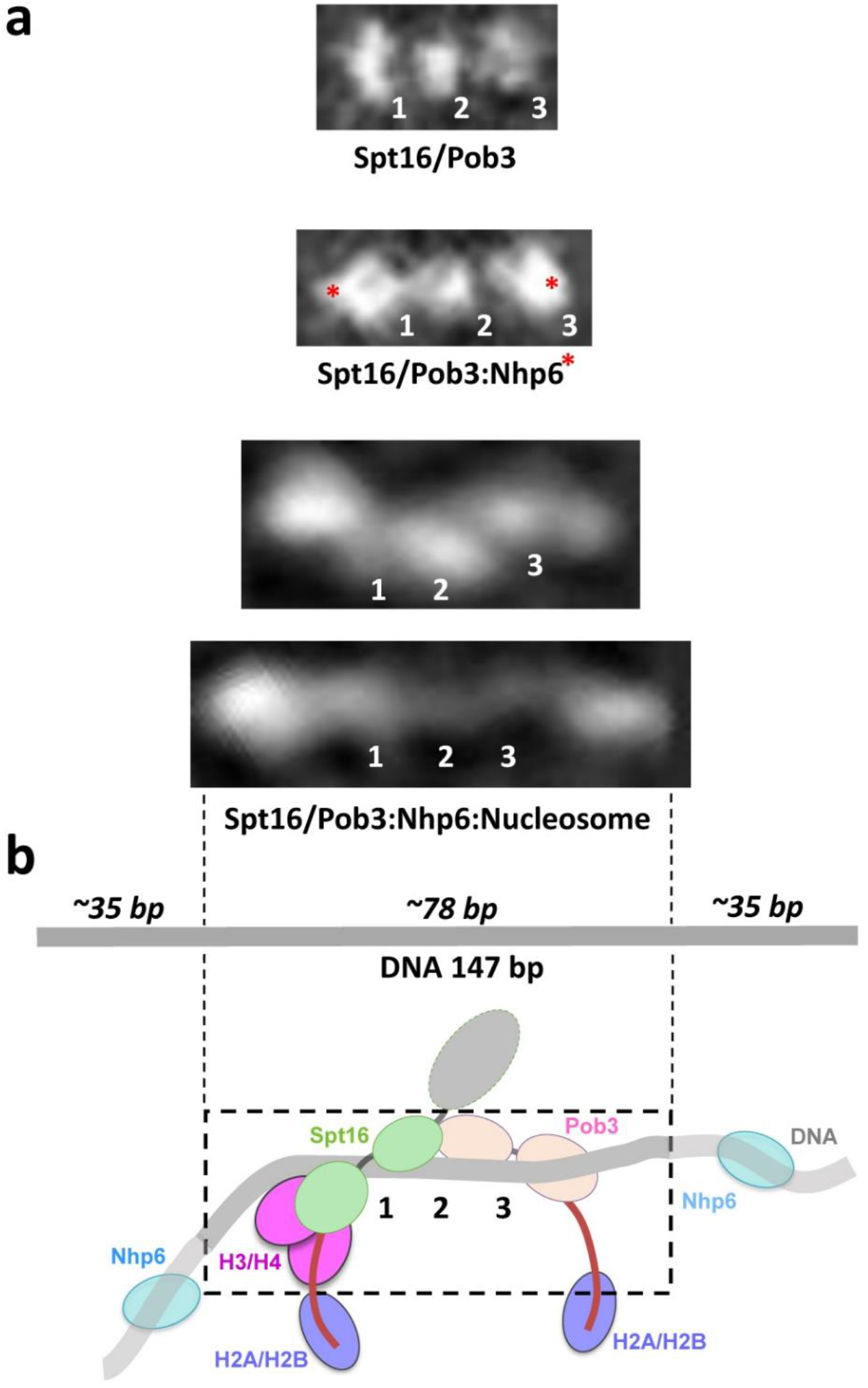
Comparison of the structures of different FACT-containing complexes. **a.** The most extended class averages for FACT, FACT:Nhp6, and FACT:Nhp6:nucleosome complexes are shown, with the densities for Nhp6 assigned from difference maps for FACT vs FACT:Nhp6 or proposed for nucleosomal complexes (red asterisks). Numbers indicate the same proposed domain assignments in each set, as in **Fig. 2b. b.** A schematic representation of the proposed domain assignments with a 147 bp DNA molecule shown to scale. The region represented in the most extended class average image is indicated by a dashed line.

In addition to the three central regions that appear to represent FACT domains (numbered 1-3 in **Fig. 4**), we also observed 2-3 additional flanking densities, with the region on the left in the orientation chosen here being larger than the one on the right (**Fig. 4a**, bottom panels). Based on the cryo-EM structure^19^, and a crystal structure of the human Spt16-M domain bound to (H3-H4)_2_ tetramers^24^, we propose that the larger density represents the Spt16-M domain bound to a histone tetramer (**Fig. 4b**). The Spt16-M domain clashes with the location of the DNA in the crystal structure, suggesting that DNA is uncoiled asymmetrically when Spt16-M binds (H3-H4)_2_. In contrast, in the form observed by cryo-EM, the dimerization domain (density 2 in our schematics) sits at the nucleosomal dyad with the M domains symmetrically positioned on either face of the nucleosome, with the DNA still coiled around the histone core^19^. We therefore propose that the dimerization domain remains associated with the DNA at the dyad as the DNA uncoils, with asymmetric extension of only the DNA that is associated with Pob3-M (**Supplementary Fig. S4**).

### A model for FACT-dependent nucleosome unfolding

Stepwise models for nucleosome unfolding by FACT:Nhp6 have been proposed, but were largely speculative. Our data support these models and extend them to incorporate a more complete set of intermediates observed by transmission EM with intact nucleosomal complexes, including steps after the uncoiling of the DNA, and proposing a new role for Nhp6 in exposing histone binding sites in FACT prior to engaging the nucleosome (**Fig. 5**). In this model, the negatively charged C-terminal tails of Spt16 and Pob3 bind to positively charged regions of the M domains, enforcing a closed conformation in which the histone-binding sites are inaccessible (**Fig. 5a**). Nhp6 first binds to the tails, promoting formation of an open structure that exposes the histone-binding sites in both M domains. Other Nhp6 molecules bind to and trap the DNA as it releases from H2A/H2B sites transiently, stabilizing exposure of the binding sites for FACT’s C-terminal tails (**Fig. 5a,b**). As H2A/H2B^22^ and Nhp6 (**Fig. 1**) both bind to the same FACT domains, we propose that this competition results in binding of the C-terminal tails to H2A/H2B and transfer of Nhp6 to the DNA. This promotes further uncoiling of the DNA, which occurs preferentially from one side of the complex due to the clash with Spt16-M, leading to asymmetrical displacement of the DNA. Multiple, incremental steps involving competing binding interactions with swapping of partners therefore lead to the formation of an extended, nearly linear structure (**Supplementary Fig. S5**), with each step facing only a small energetic barrier as each disrupted interaction is replaced quickly by a nearly equivalent one.

**Figure 5.**
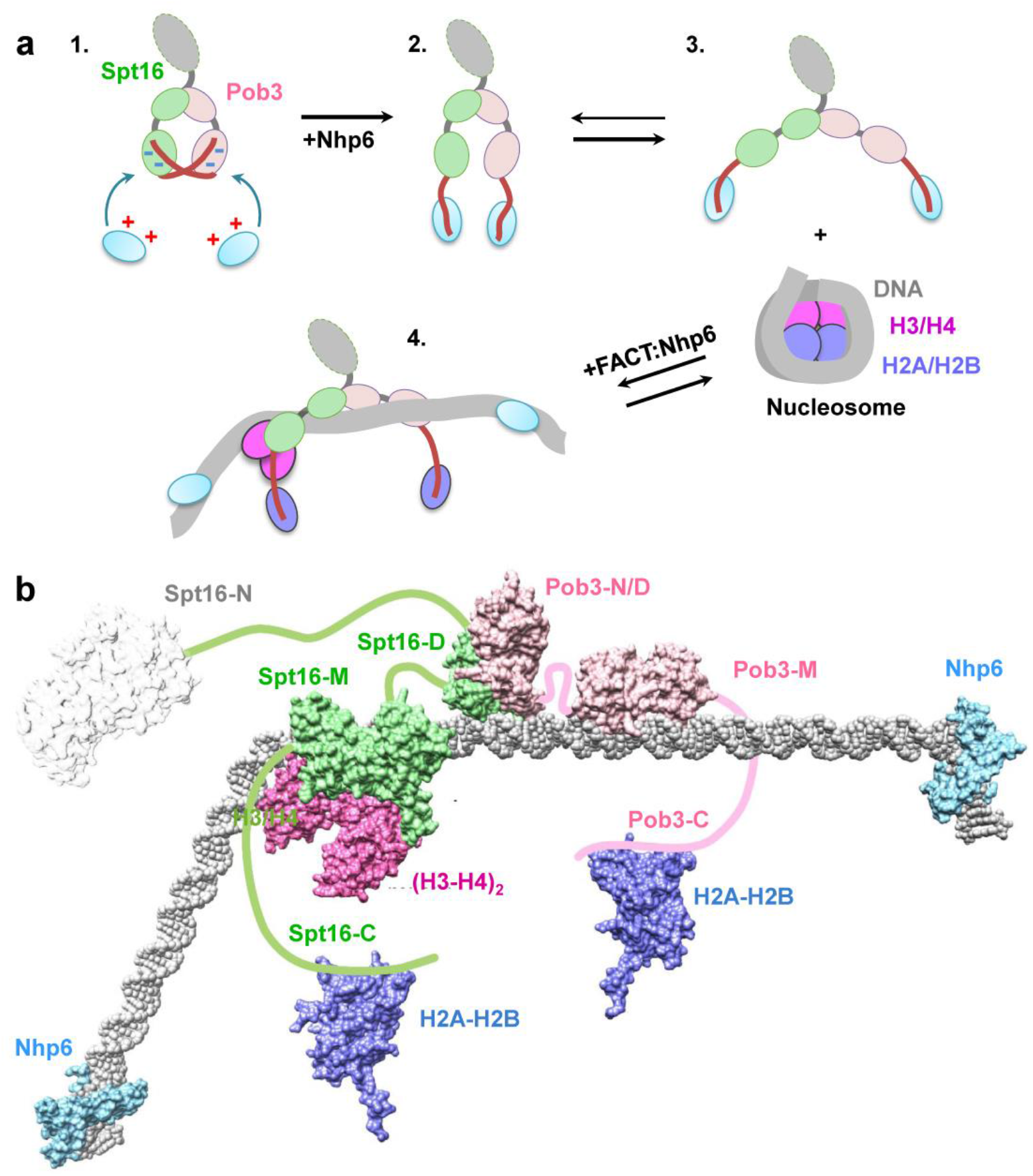
Model of nucleosome unfolding by FACT. **a.** Spt16/Pob3 is a mixture of open and closed conformations of the complex (intermediates 1, 2 and 3). Nhp6 interacts with C-terminal domains (CTDs) of Spt16 and Pob3 subunits and induces unfolding of FACT, facilitating FACT-nucleosome complex formation (intermediates 3 and 4). During nucleosome unfolding Nhp6 proteins are transferred from the CTDs to nucleosomal DNA; the vacant CTDs bind to H2A/H2B dimers that become displaced from the DNA. As a result, FACT unfolds the nucleosome in an extended, highly flexible structure (intermediate 4). Other designations as in **Fig. 2b.** The proposed structure of the unfolded FACT:Nhp6:nucleosome complex. The (H3-H4)_2_:Spt16-M:DNA complex is based on the molecular modeling described in **Supplementary Fig. S4b**, with putative positions of other components inserted using Chimera^25^ and the published structures of Spt16-N (3BIQ), Pob3-N/D:Spt16-D (4KHB), Pob3-M (2GCL), H2A/H2B (1ID3), and Nhp6:DNA (1J5N), with connectivity based on the locations of the inherently unstructured regions of each protein.

## Discussion

FACT can dramatically alter the structure of a nucleosome without ATP hydrolysis, but the extent of these changes depends on the concentration and source of HMGB-domain factors^7,17^. The SSRP1 subunit (human FACT) contains a single HMGB domain that is not found in Pob3 (yeast FACT), but both versions are capable of some activities without added factors, and both require high concentrations of the independent HMGB-family factor Nhp6 to promote full unfolding to the reorganized state^4,6,17^. We used biochemical approaches to demonstrate that Nhp6 acts on FACT in the absence of nucleosomes, and EM to show that it promotes formation of more open conformations of both FACT alone and complexes of FACT with nucleosomes. EM images revealed a broader range of structures for FACT:Nhp6:nucleosome complexes than were previously observed, supporting a stepwise series of dynamically interchangeable intermediates on a pathway from canonical nucleosomes to nearly linear, extended forms. The results answer some questions about the mechanism of nucleosome unfolding by FACT but they raise additional questions about which intermediates form under what circumstances and what their physiological roles are.

HMGB proteins bind to bent DNA^26^ so studies of the HMGB domain in SSRP1 and the separate Nhp6 protein in yeasts and fungi have focused on their potential ability to stabilize initial uncoiling of DNA at the entry/exit points of nucleosomes or to provide curvature to the DNA during nucleosome assembly. However, we found that Nhp6 can also bind to the acidic C-terminal domains of both Spt16 and Pob3 (**Fig. 1**). More importantly, this binding promoted a conformational change in FACT leading to a more open form that resembles intermediates in FACT:nucleosome complexes (**Fig. 2**). This suggests that Nhp6 is important for preparing FACT to bind to nucleosomes, possibly by exposing histone-binding sites, as well as for preparing nucleosomes to be bound by FACT, possibly by exposing histone surfaces. Initial stages of FACT binding would then involve competitions among DNA:histone, DNA:Nhp6, Nhp6:FACT, and FACT:histone interactions. Transitions among these intermediates would therefore involve little change in energy state as disrupted binding would be rapidly restored with similar interactions. Similar situations can be envisioned for subsequent steps, such as when Spt16-M engages (H3-H4)_2_ surfaces with simultaneous (asymmetrical) displacement of DNA. This raises further questions such as whether Nhp6 is transferred to the DNA during these transitions, and if so if the association is stable or dynamic.

This model also raises questions about the role of the single HMGB domain in SSRP1. The available cryo-EM structure used human FACT but the HMGB domain was not visualized^19^, suggesting it is not stably positioned in the intermediate observed. The nucleosomes in that study lacked entry/exit DNA, which in our model would make an HMGB factor unnecessary for exposing binding sites. The remaining DNA in this structure was coiled and the histone core was largely intact. The uncoiling we detect by loss of FRET in spFRET and in-gel FRET assays therefore must occur after the step represented by the cryo-EM structure, suggesting that Nhp6 is needed for this step, and that the single HMGB domain of SSRP1 is not sufficient to allow progress past this stage. What, then, is the role of this domain in SSRP1? Perhaps the abundant HMGB-family factors in mammalian cells could perform a role similar to Nhp6 in yeast.

We were able to arrange the intermediates detected by EM into a proposed sequential series of events (**Figs. 3, 5, Supplementary Fig. S5**). Our model ends in a nearly linear form consistent with the inherent rigidity of DNA, with both H2A/H2B dimers being displaced significantly. Consistently, FACT can assemble nucleosomes from core components, suggesting that it can engage histones and DNA in an even more disorganized form than detected here. The multiple structural intermediates that we did detect suggest that the range of potential configurations is larger than previously observed. To explain this range of structures, we propose that FACT populates a distributed series of multiple, energetically similar configurations of nucleosomal components. Is a similar range of different structures formed in cells and do they have different functions? For example, the extended form represented as the endpoint in our model could facilitate transcript elongation by providing a more linear DNA template, while promoting nucleosome survival by contacting each of the components of the nucleosome and tethering them together for reassembly. This or other intermediates could promote accessibility of DNA-binding proteins or enhance nucleosome eviction during promoter activation, or retention of histones during replication.

We have shown that FACT induces the formation of multiple variant nucleosomal structures and that the HMGB-family domain has several distinct roles before and during the interaction of FACT with the nucleosome. The functions of these intermediates and the distinct roles of HMGB factors in FACT function remain incompletely understood.

## Acknowledgements

Authors would like to thank Andrey Moiseenko for help with Serial EM. This work was supported by National Institutes of Health Grants R01 GM119398 (to V. M. S.) and R01 GM064649 (to T. F.). EM and spFRET studies were funded by the Russian Science Foundation (project no. 19-74-30003). EM was performed at the Unique equipment setup “3D-EMC” of Moscow Lomonosov State University. The research was partially performed using facilities of the Interdisciplinary Scientific and Educational School of Moscow University «Molecular Technologies of the Living Systems and Synthetic Biology». AKS acknowledges support by the HSE University Basic Research Program.

## Author contributions

S.A.L. performed EM and spFRET experiments, designed and performed gel-shift and native gel experiments, designed unfolding model, wrote the manuscript; M.G.K. performed EM experiments and 2D classification; M.E.V. constructed templates, designed and performed EM experiments, performed gel-shift and native gel experiments; L.L.McC. purified yFACT; T.F. designed 3D model, wrote the manuscript, interpreted data; A.K.S. designed and conducted computer modelling, wrote the manuscript; A.V.F. designed spFRET experiments, interpreted results and wrote the manuscript; M.P.K. interpreted data; S.O.S. designed experiments, performed EM and 2D classification, interpreted results and wrote the manuscript; V.M.S. designed experiments, developed unfolding model, interpreted results and wrote the manuscript.

## Supplementary Materials

**Movie S1. Flexibility of Spt16/Pob3 complex.** 2D classes of Spt16/Pob3 complexes were centered at domain 2 and are shown as a sequence of images.

**Movie S2. Flexibility of Spt16/Pob3:Nhp6 complex.** 2D classes of Spt16/Pob3:Nhp6 complex were centered at domain 2 and are shown as a sequence of images.

**Table S1. Quantitation of the open and closed FACT complexes with Nhp6 and nucleosomes.**

**Figure S1. 2D class-averages of Spt16/Pob3 (top) and Spt16/Pob3:Nhp6 complexes (bottom).**

**Figure S2. A model of FACT and Nhp6:FACT complexes in closed and open states.** Electrostatic potentials of the surfaces of relevant FACT domains and Nhp6:DNA generated in Chimera^25^ using published PDB files for Spt16-N (3BIQ), Spt16-D:Pob3-N/D (4KHB), Spt16-M (4IOY), Pob3-M (2GCL), and Nhp6:DNA (1J5N) are shown (red = −10, blue = +10 kcal/mole at 298° K). We propose that the negatively charged Spt16-C and Pob3-C domains can interact with positively charged surfaces of either FACT domains or Nhp6, swapping binding partners between the closed and open FACT states with Nhp6 binding favoring the open state.

**Figure S3. 2D Class-averages of Spt16/Pob3:Nhp6:nucleosome complexes.** Scale bar - 10 nm.

**Figure S4. Hypothetical model of a complex of tetrasome with M-domain of Spt16 subunit of yFACT. A.** H3/H4 tetrasome structure. **B**. Binding of Spt16-M domain to H3/H4 tetramer is asymmetric and incompatible with DNA binding on the surface of the tetramer^24^. To allow formation of the Spt16-M:H3/H4 tetramer complex, several DNA turns must be asymmetrically uncoiled from the surface of the tetramer.

**Figure S5. The proposed sequential pathway of nucleosome unfolding by FACT:Nhp6.** The structures of the intermediates from **Fig. 3e** were interpreted based on the assignment of the electron densities proposed in **Fig. 5b**. The resolved parts of the structures are shown by dashed lines. Other designations as in **Fig. 4b**.

## Materials and methods

### yFACT proteins

Nhp6 was expressed in *Escherichia coli* and purified as described^27,28^. WT and mutant versions of Spt16/Pob3 were purified as heterodimers from yeast cells overexpressing both proteins^29,30^. Spt16ΔC contains residues 1-958 of the 1,035 amino acid protein, and Pob3ΔC contains residues 1-477 of 552.

### Nucleosomal DNA templates

Nucleosomal DNA templates containing fluorescent labels 35 and 112 bases internal to the nucleosome boundary were amplified by PCR with the following fluorescently labeled primers: Reverse primer 5’–ACCCCAGGGACTTGAAGTAATAAGGACGGAGGGCC**T#**CTTTCAACATCGAT (where T# - is a nucleotide labeled with Cy3), Forward primer 5’–CCCGGTTCGCGCTCCCT CCTTCCGTGTGTTGTCG**T\***CTCT (where T* - is a nucleotide labeled with Cy5).

A plasmid containing the modified Widom 603–42 sequence^31^ was purified with a QIAquick PCR Purification Kit (Qiagen) and used as the template for the amplification.

### Nucleosome assembly and purification

Recombinant histones from *Xenopus laevis* were expressed in *Escherichia coli* and purified as described^32^. Nucleosomes were assembled with recombinant octamers by dialysis from 2 M NaCl as described^33,34^.

### EMSA of FACT-nucleosome complexes

Formation of FACT complexes with nucleosomes was evaluated using an electrophoretic mobility shift assay (EMSA) as described^7,8,10^ after incubation in a buffer A containing 17 mM HEPES pH 7.6, 2 mM Tris-HCl, 0.8 mM Na_3_EDTA, 0.11 mM β-mercaptoethanol, 11 mM NaCl, 1.1% glycerin, and 12% sucrose.

Spt16/Pob3 was used at a final concentration of 0.13 μM and Nhp6 at a final concentration of 1.3 μM. Nucleosomes were added to a final concentration of ~10 nM for detecting in EMSA analysis and ~ 30 nM for EM. Intact nucleosomes and FACT:Nhp6 complexes were detected by in-gel FRET. The gel was scanned using a Typhoon scanner (GE Healthcare, USA) with excitation at 532 nm laser and emission at 670 nm (Cy3-Cy5 FRET) or 580 nm (Cy3 signal) as described^35^.

### Analysis of the protein content of FACT:Nhp6 and FACT:Nhp6:nucleosome complexes by 2-dimensional gel electrophoresis

Complexes were formed by incubating Spt16/Pob3 (0.13 μM) and Nhp6 (1.3 μM) in buffer A for 10 min at 30 °C and separated by native PAGE as described above using 4% PAAG (AA:Bis =39:1) at 4 °C. The bands containing the complexes were excised, then crushed and incubated for 15 h at 4 °C with an equal volume of HE buffer (10 mM HEPES-NaOH, pH 8.0, 0.2 mM EDTA), then finally washed with 50 – 100 μL of additional HE buffer. The supernatant containing the proteins was recovered after centrifugation and mixed with 4X buffer (200 mM Tris-HCl pH 6.8, 400 mM β-mercaptoethanol, 4% sodium dodecyl sulfate (SDS), 40% glycerol). After incubating at 95 °C for 5 min with periodic vortexing, samples were separated on an Invitrogen Bolt 4-12% Bis-Tris gel (Invitrogen, USA) with MES SDS Running buffer (ThermoFisher Scientific, USA), and proteins were detected by silver staining (SilverQuest Staining Kit, Invitrogen, USA).

### Preparation of samples for electron microscopy: FACT and FACT:Nhp6

Spt16/Pob3 samples with or without Nhp6 were analyzed in buffer A as for the EMSA assay^7^. Spt16/Pob3 and Nhp6 were used at final concentrations of 0.13 and 1.3 μM, respectively. Reaction mixtures were incubated for 10 min at 30 °C, then 3 μl samples (0.133 μM complexes) were place on copper grids (300 mesh formvar/carbon-coated) (Ted Pella, USA) that were hydrophilized by glow discharge (−20 mA, 45 s) with an Emitech K100X. Grids were negatively stained with 1% uranyl acetate (Spi, USA).

### Preparation of samples for electron microscopy: FACT:Nhp6:nucleosome

FACT:Nhp6:nucleosome complexes were formed by incubating Spt16/Pob3 (0.13 μM), Nhp6 (1.3 μM) and nucleosomes (10 nM) for 10 min at 30°C in buffer A. Complexes were purified by native PAGE (100 V for ~50 min in 0.5X TBE), then the Cy3/Cy5 labels were detected with the Typhoon scanner. The gel was placed into a humidified chamber, and the band of interest was excised. Carbon-coated copper grids (SPI, United States) were glow discharged for 2 min as described above and immediately placed with the charged side down on the scratched surface of the native gel in 0.5X TBE buffer and incubated for 5 min. Excess buffer was removed from the grid and the grid was subjected to negative contrasting with a 1% solution of uranyl acetate for 30 s at 25 °C.

### Electron microscopy and image analysis

Samples were analyzed using a Jem 2100 analytical transmission electron microscope (JEOL, Japan) equipped with a 2K × 2K CCD camera Ultrascan 1000XP (Gatan, USA). The microscope was operated at 200kV, with a magnification of 40,000x (2.5 Å/pix) and a defocus of 0.5–1.9 μm. Images were acquired with SerialEM software in the low dose mode^36^.

Micrographs were imported to the Eman2 suite^37,38^ and CTF-corrected. A training subset of individual particles was selected manually with EMAN2 boxer, then others were acquired automatically using the crYOLO neural network^39^. Box coordinates were imported to Eman2 where particles were subjected to alignment and 2D classification.

Particles of FACT-nucleosome complexes were exported to Relion2.0.5^40^ for 2D classification and analysis. Difference maps and measuring of the 2D class dimensions were performed in Fiji^41^.

### spFRET experiments

Nucleosomes containing fluorescent labels at positions 35 and 112 bases internal to the nucleosome boundary were gel purified and used for spFRET measurements at a concentration of 0.5 nM as described^7^. Nucleosomes were incubated in the presence of Spt16/Pob3 (0.13 μM) and Nhp6 (1.3 μM) for 10 min at 30°C in buffer A.

Three independent series of experiments were performed, and the data (1600-8800 signals for each measured sample) were presented as relative frequency distributions of nucleosomes by proximity ratio E_PR_ (E_PR_ profiles) as described^7,42^. E_PR_ profiles were further approximated as a superposition of two Gaussian curves, where each Gaussian corresponded to a particular subpopulation of nucleosomes with different FRET profile. The content of each nucleosome subpopulation was calculated as the ratio of the area under the corresponding Gaussian peak to the area under the entire E_PR_-profile. The E_PR_ profiles and contents of nucleosome subpopulations were averaged (mean±SEM) over three independent experiments.

### Modeling a complex of tetrasome with M-domain of Spt16 subunit of yFACT

Tetrasome structure was prepared from nucleosome core particle structure (PDB ID 1KX5^43^) by removing H2A-H2B histone dimers and uncoiling 53 bp of DNA from each nucleosomal DNA end. Next a structure of histone H3-H4 tetramer with Spt16 M-domain (PDB ID 4Z2M^24^) was superimposed onto the tetrasome structure. Clashes between Spt16 and DNA were monitored with UCSF Chimera^25^. 80 DNA bp from one nucleosomal DNA end had to be uncoiled to avoid clashes. For DNA uncoiling 3DNA software suite was used^44^ which allowed to rebuild parts of DNA structure with base pair and base pair step parameters set to those corresponding to the straight canonical B-DNA.

## References and Notes

1. Luger, K., Mader, A.W., Richmond, R.K., Sargent, D.F. & Richmond, T.J. Crystal structure of the nucleosome core particle at 2.8 A resolution. Nature 389, 251–60 (1997).

2. Vasudevan, D., Chua, E.Y.D. & Davey, C.A. Crystal structures of nucleosome core particles containing the ‘601’ strong positioning sequence. J Mol Biol 403, 1–10 (2010).

3. Clapier, C.R., Iwasa, J., Cairns, B.R. & Peterson, C.L. Mechanisms of action and regulation of ATP-dependent chromatin-remodelling complexes. Nat Rev Mol Cell Biol 18, 407–422 (2017).

4. Formosa, T. & Winston, F. The role of FACT in managing chromatin: disruption, assembly, or repair? Nucleic Acids Res 48, 11929–11941 (2020).

5. Zhou, K., Liu, Y. & Luger, K. Histone chaperone FACT FAcilitates Chromatin Transcription: mechanistic and structural insights. Curr Opin Struct Biol 65, 26–32 (2020).

6. Gurova, K., Chang, H.W., Valieva, M.E., Sandlesh, P. & Studitsky, V.M. Structure and function of the histone chaperone FACT - Resolving FACTual issues. Biochim Biophys Acta Gene Regul Mech 1861, 892–904 (2018).

7. Valieva, M.E. et al. Large-scale ATP-independent nucleosome unfolding by a histone chaperone. Nat Struct Mol Biol 23, 1111–1116 (2016).

8. Xin, H. et al. yFACT induces global accessibility of nucleosomal DNA without H2A-H2B displacement. Mol Cell 35, 365–76 (2009).

9. Formosa, T. et al. Spt16-Pob3 and the HMG protein Nhp6 combine to form the nucleosome-binding factor SPN. EMBO J 20, 3506–17 (2001).

10. Valieva, M.E. et al. Stabilization of Nucleosomes by Histone Tails and by FACT Revealed by spFRET Microscopy. Cancers (Basel) 9, pii: E3 (2017).

11. Erkina, T.Y. & Erkine, A. ASF1 and the SWI/SNF complex interact functionally during nucleosome displacement, while FACT is required for nucleosome reassembly at yeast heat shock gene promoters during sustained stress. Cell Stress Chaperones 20, 355–69 (2015).

12. Takahata, S., Yu, Y. & Stillman, D.J. FACT and Asf1 regulate nucleosome dynamics and coactivator binding at the HO promoter. Mol Cell 34, 405–15 (2009).

13. Hsieh, F.K. et al. Histone chaperone FACT action during transcription through chromatin by RNA polymerase II. Proc Natl Acad Sci U S A 110, 7654–9 (2013).

14. Cheung, V. et al. Chromatin- and transcription-related factors repress transcription from within coding regions throughout the *Saccharomyces cerevisiae* genome. PLoS Biol 6, e277 (2008).

15. Jamai, A., Imoberdorf, R.M. & Strubin, M. Continuous histone H2B and transcription-dependent histone H3 exchange in yeast cells outside of replication. Mol Cell 25, 345–55 (2007).

16. Voth, W.P. et al. A role for FACT in repopulation of nucleosomes at inducible genes. PLoS One 9, e84092 (2014).

17. McCullough, L.L. et al. Functional roles of the DNA-binding HMGB domain in the histone chaperone FACT in nucleosome reorganization. J Biol Chem 293, 6121–6133 (2018).

18. Chang, H.W. et al. Mechanism of FACT removal from transcribed genes by anticancer drugs curaxins. Sci Adv 4, eaav2131 (2018).

19. Liu, Y. et al. FACT caught in the act of manipulating the nucleosome. Nature 577, 426–431 (2020).

20. Brewster, N.K., Johnston, G.C. & Singer, R.A. A bipartite yeast SSRP1 analog comprised of Pob3 and Nhp6 proteins modulates transcription. Mol Cell Biol 21, 3491–502 (2001).

21. Stros, M. HMGB proteins: interactions with DNA and chromatin. Biochim Biophys Acta 1799, 101–13 (2010).

22. Kemble, D.J., McCullough, L.L., Whitby, F.G., Formosa, T. & Hill, C.P. FACT Disrupts Nucleosome Structure by Binding H2A-H2B with Conserved Peptide Motifs. Mol Cell 60, 294–306 (2015).

23. Thastrom, A. et al. Sequence motifs and free energies of selected natural and non-natural nucleosome positioning DNA sequences. J Mol Biol 288, 213–29 (1999).

24. Tsunaka, Y., Fujiwara, Y., Oyama, T., Hirose, S. & Morikawa, K. Integrated molecular mechanism directing nucleosome reorganization by human FACT. Genes Dev 30, 673–86 (2016).

25. Pettersen, E.F. et al. UCSF Chimera--a visualization system for exploratory research and analysis. J Comput Chem 25, 1605–12 (2004).

26. Masse, J.E. et al. The S. cerevisiae architectural HMGB protein NHP6A complexed with DNA: DNA and protein conformational changes upon binding. J Mol Biol 323, 263–84 (2002).

27. Ruone, S., Rhoades, A.R. & Formosa, T. Multiple Nhp6 molecules are required to recruit Spt16-Pob3 to form yFACT complexes and to reorganize nucleosomes. J Biol Chem 278, 45288–95 (2003).

28. Paull, T.T. & Johnson, R.C. DNA looping by Saccharomyces cerevisiae high mobility group proteins NHP6A/B. Consequences for nucleoprotein complex assembly and chromatin condensation. J Biol Chem 270, 8744–54 (1995).

29. Biswas, D., Yu, Y., Prall, M., Formosa, T. & Stillman, D.J. The yeast FACT complex has a role in transcriptional initiation. Mol Cell Biol 25, 5812–22 (2005).

30. Wittmeyer, J., Joss, L. & Formosa, T. Spt16 and Pob3 of Saccharomyces cerevisiae form an essential, abundant heterodimer that is nuclear, chromatin-associated, and copurifies with DNA polymerase alpha. Biochemistry 38, 8961–71 (1999).

31. Kulaeva, O.I. et al. Mechanism of chromatin remodeling and recovery during passage of RNA polymerase II. Nat Struct Mol Biol 16, 1272–8 (2009).

32. Luger, K., Rechsteiner, T.J., Flaus, A.J., Waye, M.M. & Richmond, T.J. Characterization of nucleosome core particles containing histone proteins made in bacteria. J Mol Biol 272, 301–11 (1997).

33. Kireeva, M.L. et al. Nucleosome remodeling induced by RNA polymerase II: loss of the H2A/H2B dimer during transcription. Mol Cell 9, 541–52 (2002).

34. Gaykalova, D.A., Kulaeva, O.I., Bondarenko, V.A. & Studitsky, V.M. Preparation and analysis of uniquely positioned mononucleosomes. Methods Mol Biol 523, 109–23 (2009).

35. Sultanov, D.C. et al. Unfolding of core nucleosomes by PARP-1 revealed by spFRET microscopy. AIMS Genet 4, 21–31 (2017).

36. Mastronarde, D.N. Automated electron microscope tomography using robust prediction of specimen movements. J Struct Biol 152, 36–51 (2005).

37. Tang, G. et al. EMAN2: an extensible image processing suite for electron microscopy. J Struct Biol 157, 38–46 (2007).

38. Bell, J.M., Chen, M., Baldwin, P.R. & Ludtke, S.J. High resolution single particle refinement in EMAN2.1. Methods 100, 25–34 (2016).

39. Wagner, T. et al. SPHIRE-crYOLO is a fast and accurate fully automated particle picker for cryo-EM. Commun Biol 2, 218 (2019).

40. Scheres, S.H. et al. Maximum-likelihood multi-reference refinement for electron microscopy images. J Mol Biol 348, 139–49 (2005).

41. Schindelin, J. et al. Fiji: an open-source platform for biological-image analysis. Nat Methods 9, 676–82 (2012).

42. Kudryashova, K.S. et al. Preparation of mononucleosomal templates for analysis of transcription with RNA polymerase using spFRET. Methods Mol Biol 1288, 395–412 (2015).

43. Davey, C.A., Sargent, D.F., Luger, K., Maeder, A.W. & Richmond, T.J. Solvent mediated interactions in the structure of the nucleosome core particle at 1.9 a resolution. J Mol Biol 319, 1097–113 (2002).

44. Lu, X.J. & Olson, W.K. 3DNA: a versatile, integrated software system for the analysis, rebuilding and visualization of three-dimensional nucleic-acid structures. Nat Protoc 3, 1213–27 (2008).

