## Supplemental materials for "ATP-Independent Nucleosome Unfolding by FACT: Electron Microscopy Analysis"

### Supplemental Data

**Table S1. Quantitation of the open and closed FACT complexes with Nhp6 and nucleosomes.**

**Figure S1. 2D class-averages of Spt16/Pob3 (top) and Spt16/Pob3:Nhp6 complexes (bottom).**

**Figure S2. A model of FACT and Nhp6:FACT complexes in closed and open states.**

**Figure S3. 2D Class-averages of Spt16/Pob3:Nhp6:nucleosome complexes. Scale bar - 10 nm.**

**Figure S4. Hypothetical model of a complex of tetrasome with M-domain of Spt16 subunit of yFACT.**

**Figure S5. The proposed sequential pathway of nucleosome unfolding by FACT:Nhp6.**

**Table S1. Quantitation of the open and closed FACT complexes with Nhp6 and nucleosomes.** .

| Sample | N particles | N particles ≤90° (closed) | % ≤90° (closed) | N particles >90° (open) | % >90° (open) |
|---|---|---|---|---|---|
| **Spt16/Pob3** | 10304 | 6594 | 64,0 | 3710 | 36,0 |
| **Spt16/Pob3 + Nhp6** | 28425 | 13829 | 48,7 | 14596 | 51,3 |
| **N + Spt16/Pob3 + Nhp6** | 2139 | 963 | 45,0 | 1176 | 55,0 |
| **N + Nhp6** | 3251 | 3251 | 100,0 | 0 | 0,0 |

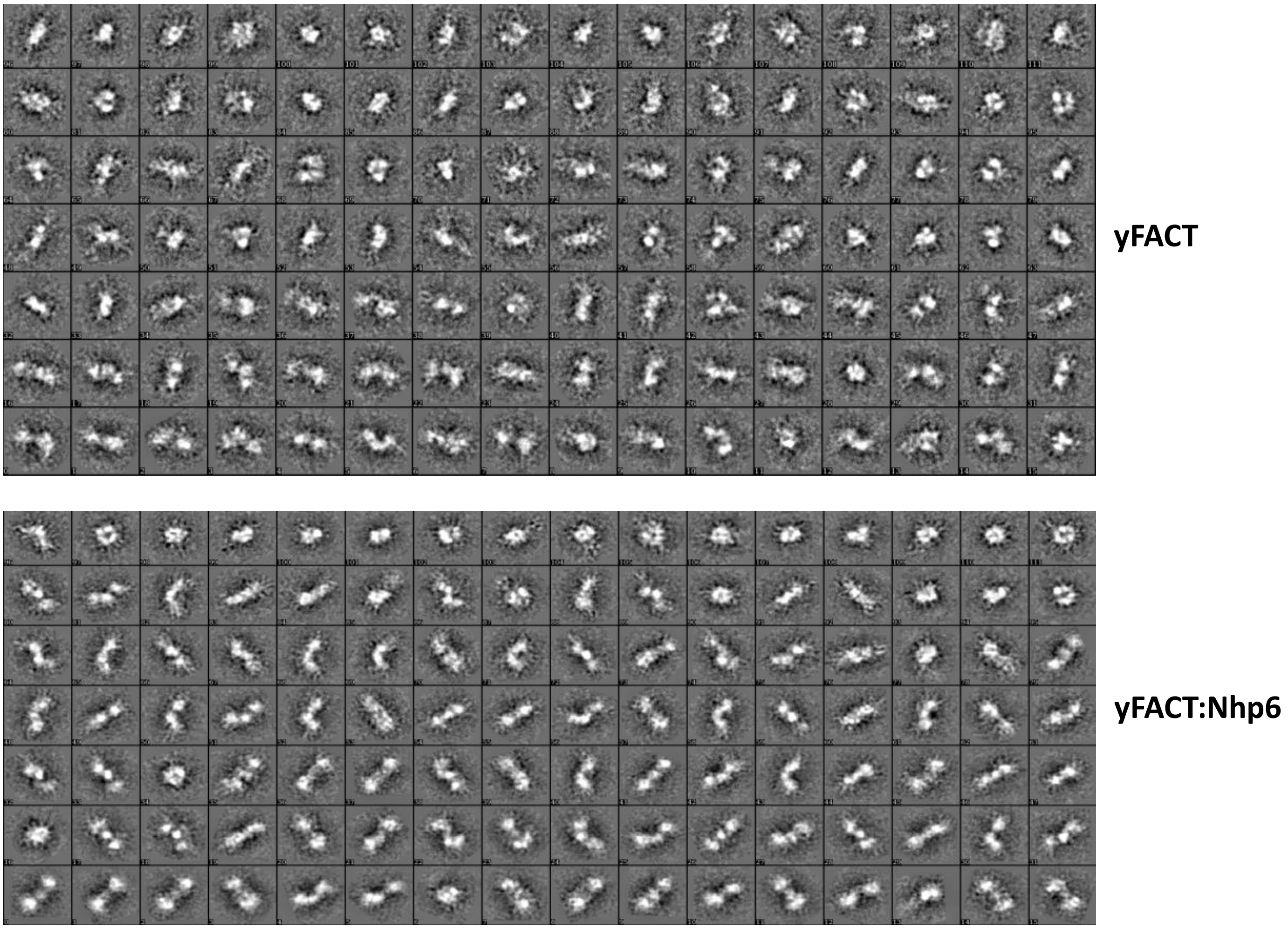


**Figure S1. 2D class-averages of Spt16/Pob3 (top) and Spt16/Pob3:Nhp6 complexes (bottom).**

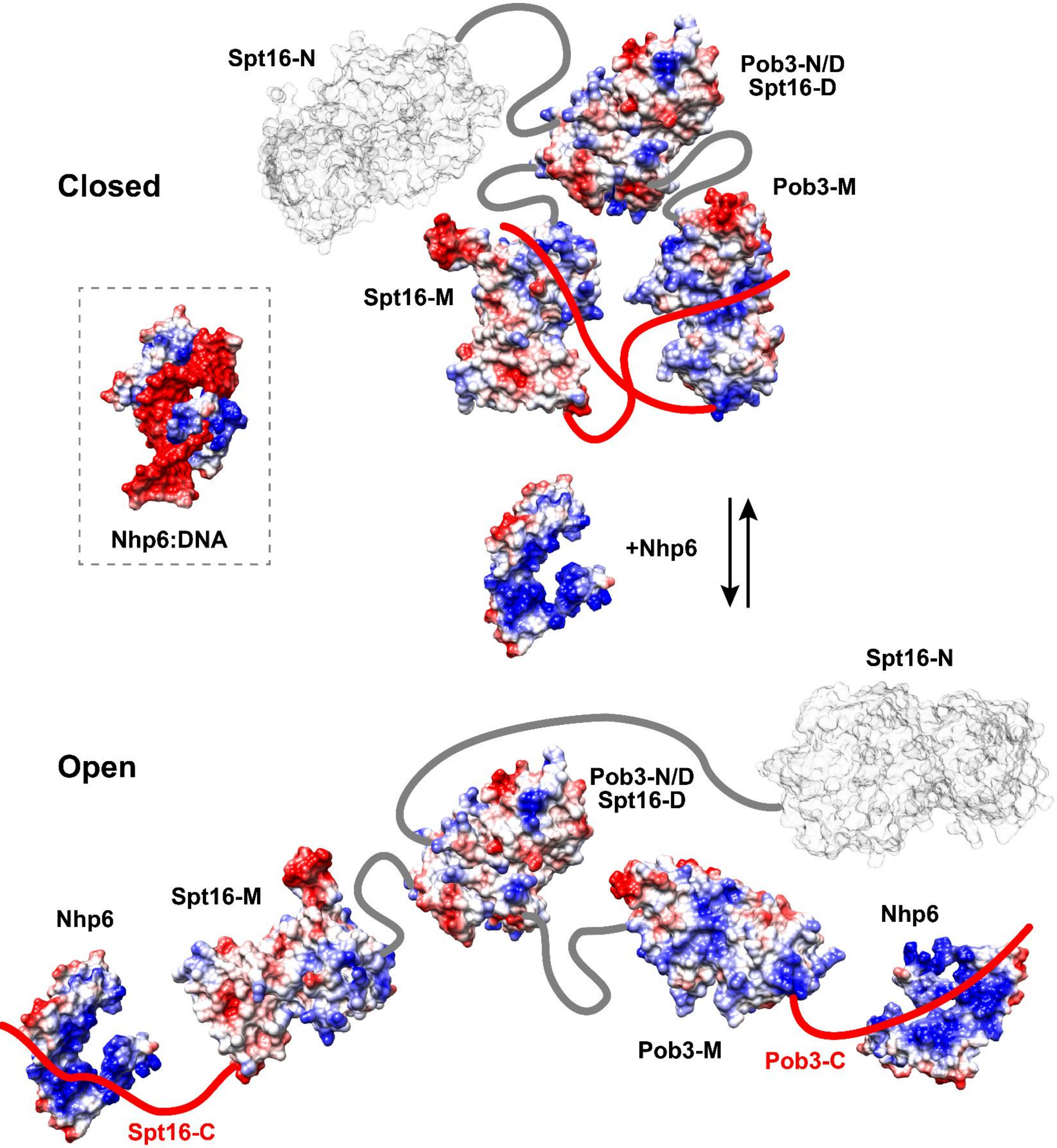


**Figure S2. A model of FACT and Nhp6:FACT complexes in closed and open states.** Electrostatic potentials of the surfaces of relevant FACT domains and Nhp6:DNA generated in Chimera[25] using published PDB files for Spt16-N (3BIQ), Spt16-D:Pob3-N/D (4KHB), Spt16-M (4IOY), Pob3-M (2GCL), and Nhp6:DNA (1J5N) are shown (red = -10, blue = +10 kcal/mole at 298° K). We propose that the negatively charged Spt16-C and Pob3-C domains can interact with positively charged surfaces of either FACT domains or Nhp6, swapping binding partners between the closed and open FACT states with Nhp6 binding favoring the open state.

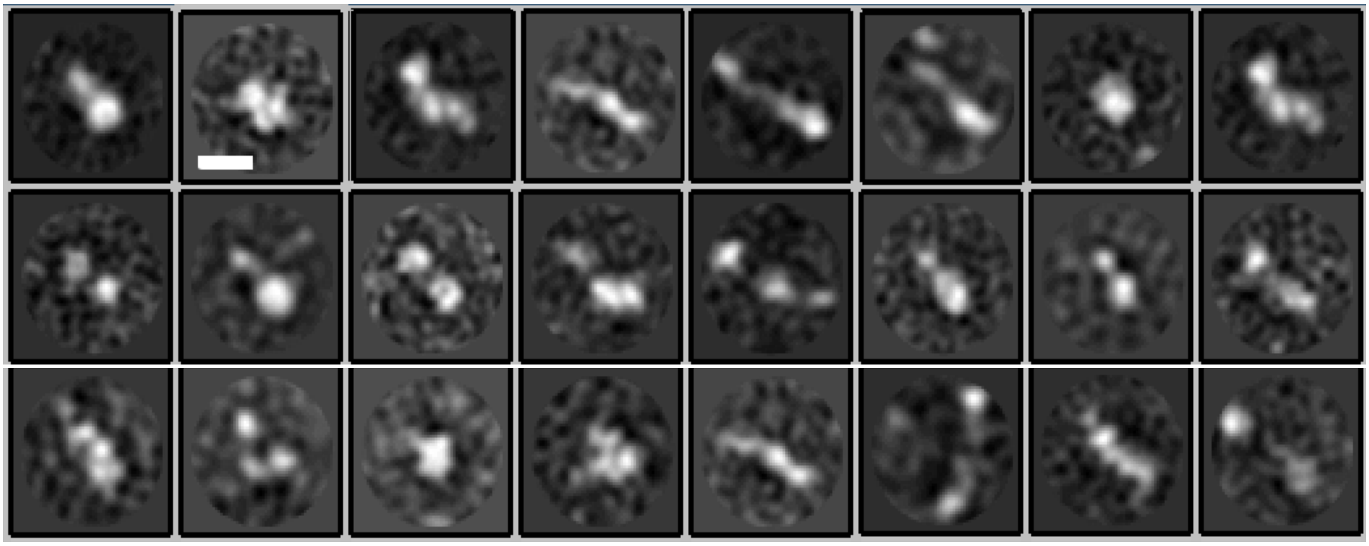

**Figure S3. 2D Class-averages of Spt16/Pob3:Nhp6:nucleosome complexes.** Scale bar – 10 nm.

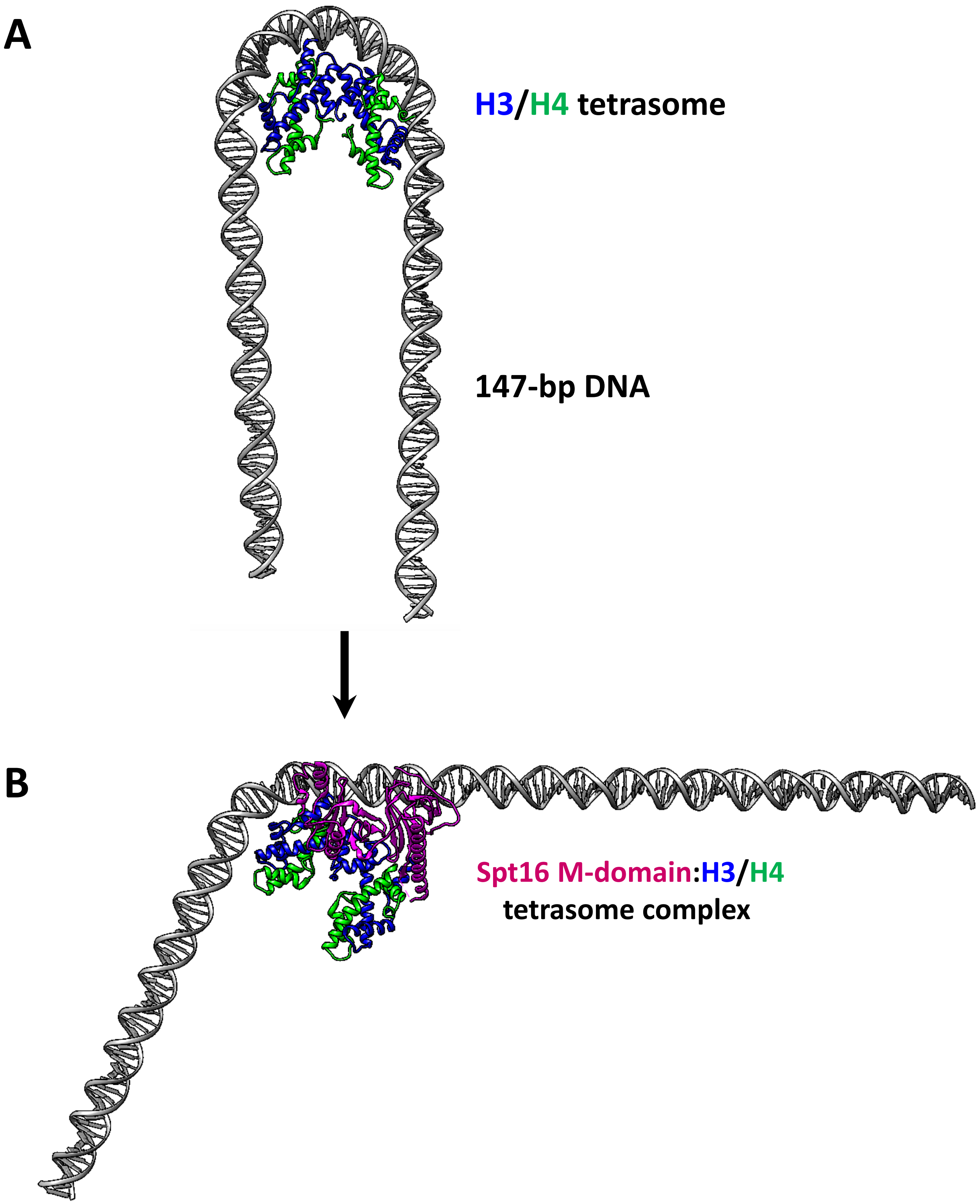


**Figure S4. Hypothetical model of a complex of tetrasome with M-domain of Spt16 subunit of yFACT. A.** H3/H4 tetrasome structure. **B.** Binding of Spt16-M domain to H3/H4 tetramer is asymmetric and incompatible with DNA binding on the surface of the tetramer. To allow formation of the Spt16-M:H3/H4 tetramer complex, several DNA turns must be asymmetrically uncoiled from the surface of the tetramer.

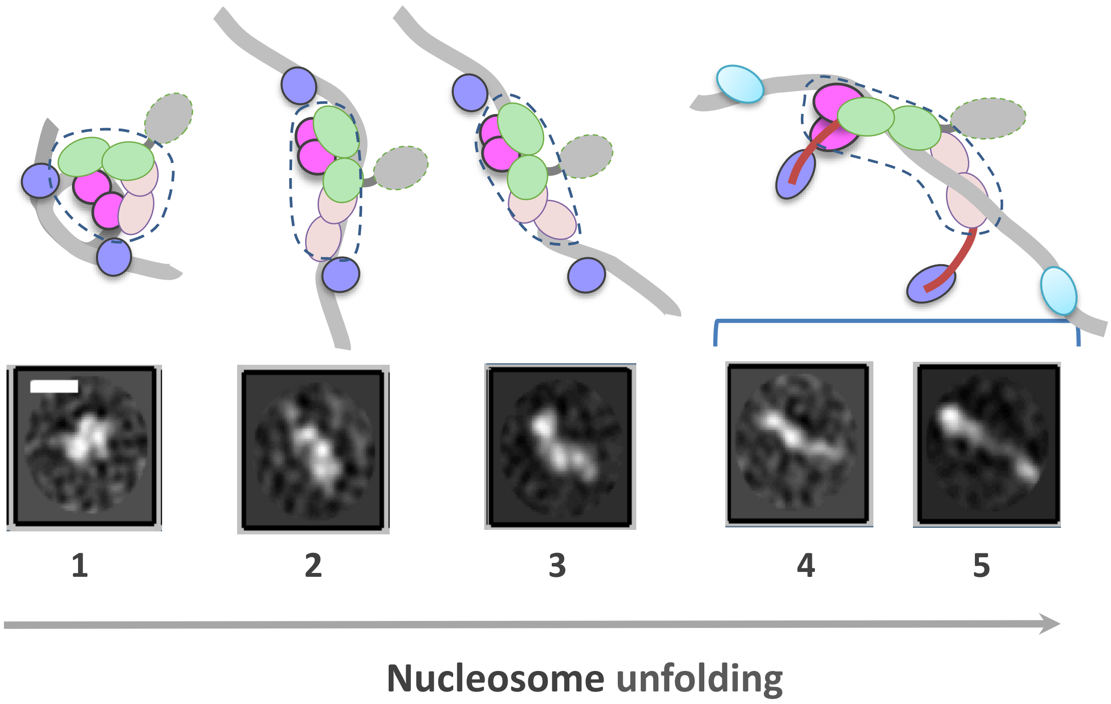


**Figure S5. The proposed sequential pathway of nucleosome unfolding by FACT:Nhp6.** The structures of the intermediates from **Fig. 3E** were interpreted based on the assignment of the electron densities proposed in **Fig. 5B**. The resolved parts of the structures are shown by dashed lines. Other designations as in **Fig. 4B**.
